# *Eigengene* reveals invariant global spatial patterns across mouse and fish brain development

**DOI:** 10.1101/2024.09.19.613507

**Authors:** Stan Kerstjens, Florian Engert, Anthony M Zador, Rodney J Douglas

**Affiliations:** Cold Spring Harbor Laboratory; Department of Molecular and Cellular Biology, Harvard University; Institute of Neuroinformatics, UZH & ETH Zurich

## Abstract

Development from a zygote to an adult organism involves complex interactions among thousands of genes. These genes exhibit highly dynamic expression across space and time. Here we report a striking simplicity amidst this complexity: Despite individual gene expression variability, the *eigengene*—the principal component of gene expression—exhibits an invariant global spatial pattern throughout the embryonic and post-natal stages of the mouse brain. Furthermore, the mouse pattern is observed also in the larval zebrafish, revealing that eigengene expression is conserved over 400 million years of evolution. We show that the eigengene pattern can be explained by a simple lineage model in which daughter cells’ gene expression is similar to that of their parent, but cannot be explained by one in which gene expression arises through local cellular signaling. The constrained lineage gives rise naturally to a global eigengene expression hierarchy that could aid in the formation of a spatial hierarchy of long-range signal gradients. We propose that lineage thus induces an address-like organization, which could have been co-opted by evolution for developmental processes that require positional information over a wide range of spatial scales, such as tissue patterning and axon navigation.

---

The development of an organism from an initial zygote unfolds through a sequence of developmental events, regulated by the complex interactions among thousands of genes. This highly stereotyped process ultimately gives rise to the diverse array of cell types and tissues that make up the mature organism. One of the most striking features of this process is the highly dynamic and tightly regulated nature of gene expression: Individual genes are turned on and off with exquisite spatial and temporal specificity, with each cell expressing a specific combination of genes at each developmental stage (Davidson & Peter, 2015). It is these complex stereotyped patterns that sketch out the detailed body plan, and provide individual cells with the positional information they need to choose their proper fates, migrate, and navigate their many processes such as neuronal axons (Stoeckli, 2018; Wolpert, 1969). Deciphering the principles that govern these expression patterns is key to understanding the nature of organismal organization, and the developmental process that brings it into being (Davidson & Peter, 2015; Waddington, 1956).

Establishing the spatio-temporal patterning of tissue represents a challenging computational process. It was indeed Alan Turing, one of the fathers of computing, who formulated the first mathematical models for how the simple interactions among a few key chemicals, for which he coined the term *morphogens*, can generate complex spatial expression patterns in developing tissue (Turing, 1952). This and similar models show how simple interactions among few genes can generate complex spatio-temporal patterns (Gierer & Meinhardt, 1972; Green & Sharpe, 2015; Jaeger, 2011; Mandelbrot, 1983). However, the converse is also true: Seemingly simple patterns, such as long-range concentration gradients, can be surprisingly complex to generate. Lewis Wolpert and Francis Crick were among the first to recognize that although diffusion is an effective method for establishing gradients over relatively short distances (50–100 cells), physical constraints imposed by diffusion prevent such simple mechanisms from scaling to larger distances (Crick, 1970; Francis & Palsson, 1997; Goodhill, 2016; Wolpert, 1969) and instead require increasingly complex regulatory logic and machinery such as active transport, relays, and extracellular binders (Briscoe & Small, 2015; Stapornwongkul & Vincent, 2021).

How developmental processes orchestrate and convey positional information across long spatial scales through cellular-level gene regulation remains unclear. This largescale positional information plays a crucial role in guiding processes that depend on global cues, even in later developmental stages, such as cell migration and axon navigation. The challenge is further complicated by the genome’s limited capacity to encode extensive regulatory logic (Kerstjens et al., 2022; Zador, 2019). This constraint renders ineffective certain strategies that, while potentially viable on a local scale, would require too much time or regulatory logic to scale globally. These include strategies that require axons to exhaustively search large regions of tissue for their synaptic partners, or that require large numbers of cells to be tagged with unique yet specific chemical labels (Goodhill & Xu, 2005; Sperry, 1963; von der Malsburg, 1987).

Here, we use brain-wide maps of gene expression, along with novel analyses, to uncover a surprisingly ubiquitous system of global gene expression patterns. We report that patterns of gene expression are remarkably conserved across ontogeny and phylogeny, even in the face of highly variable changes in the expression of individual genes. Using principal component analysis (PCA), we identify a stable feature of gene co-expression patterns, the *principal eigengene expression*. Strikingly, the spatial expression of the principal eigengene remains largely unchanged during the brain development of mice, and is even conserved across zebrafish. We demonstrate that such stable gene expression can be induced by intrinsic constraints of the cell division process, and so does not necessarily require complex gene regulatory machinery. Our lineal model predicts global eigengene expression patterns that persist across the tissue throughout development, forming a global eigengene hierarchy. In contrast, a purely neighborhood model, relying solely on local cell-cell interactions, fails to produce such spatially scalable and temporally persistent patterns. These findings provide a framework for understanding the local coordinate spaces built on local molecular gradients, suggesting that they may be part of an overarching global hierarchy of eigengene expression gradients that naturally arise from the cell division process. This global eigengene hierarchy offers a simple framework for understanding the coordination of seemingly independent local gradients across time and space, and may play a crucial role in guiding large-scale developmental processes by providing global positional information.

## Results

We present five results. First, spatial eigengene expression forms a static global pattern throughout development, despite the fact that individual gene expression is highly variable (Thompson et al., 2014). Second, the eigengene measured in the mouse brain has a surprisingly similar spatial expression pattern in the larval zebrafish brain, despite a separation of more than 400 million years of evolution between the two species. Third, any random subset of about 50 genes shows largely the same spatial pattern in both mouse and zebrafish. Fourth, a simple lineal model, in which gene expression of daughter cells is similar to that of their parent, can induce the observed eigengene pattern. This lineal model explains the pattern’s global and persistent nature, despite the highly dynamic nature of individual genes. Furthermore, the model predicts that eigengenes organize the tissue into a hierarchy of contiguous regions. Finally, the predicted hierarchical regions indeed exist, and are contiguous, in both mouse and zebrafish data.

### Individual genes fluctuate across development

The gene expression data we analyzed are drawn from previously published studies, including the Allen Institute (Thompson et al., 2014) and Mapzebrain (Shainer et al., 2023). They consist of voxels of gene expression measured at 4 embryonic stages (E11.5, E13.5, E15.5, and E18.5) and 4 postnatal stages (P4, P14, P28, and P56) in mouse (Thompson et al., 2014), and one larval stage (6dpf) in zebrafish (Shainer et al., 2023). Each mouse voxel contains the expression values of 1256 genes, and each zebrafish voxel of 290 genes. Of these genes, 45 homologues are present in both data sets. In the case of the mouse, the voxel sizes range from 80 to 200 microns according to the embryonic stage at which the data was measured (Thompson et al., 2014). For the zebrafish the voxels are much smaller, approaching singlecell diameter (Shainer et al., 2023).

Spatial gene expression is highly dynamic over developmental time. Fig. 1a shows the spatial expression of 5 randomly selected genes across the measured stages of mouse brain development, and one stage of the larval zebrafish brain. These representative examples indicate that spatial gene expression often undergoes considerable changes over the course of development (Fig. 1b). Indeed, when all individual voxels of the mouse brain atlas are joined in one T-SNE embedding space, the voxels cluster with other voxels measured at the same developmental stage, rather than with voxels of the same tissue at different stages (Fig. 1c). Moreover, not only does spatial pattern of expression change over the course of development, but the correlational structure among genes also changes (Fig. 2d). These analyses reinforce the view that at the level of single genes and correlations, gene expression is highly dynamic across development.

**Figure 1:**
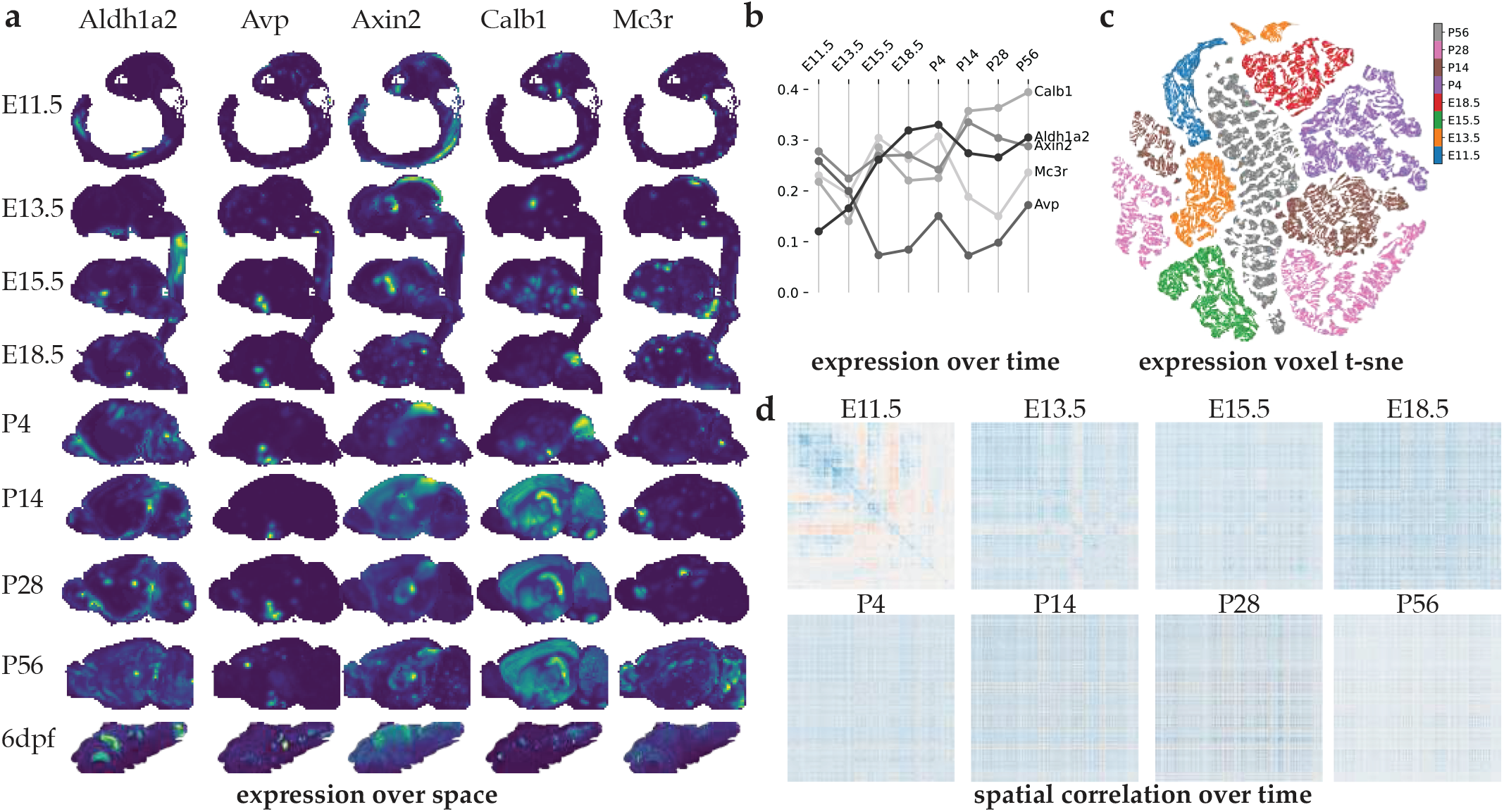
Expression of single genes is spatially heterogeneous and fluctuates across development. (**a**) Voxels of gene expression were measured through *in situ* hybridization at 4 embryonic stages and 4 postnatal stages in mouse (Thompson et al., 2014), and one larval stage (6pdf) in zebrafish (Shainer et al., 2023). Each mouse voxel contains the expression of 1256 genes, and each zebrafish voxel 290 genes. Of these genes, 45 homologues are present in both data sets. The spatial expression of five representative (randomly selected) genes varies through development. (**b**) The averaged and normalized temporal expression of the same five genes throughout mouse development. (**c**) A T-SNE embedding of all voxels in the mouse data. Voxels are colored by their developmental time point. (**d**) The correlation matrices among all genes in the mouse expression data. Blue and red indicate negative and positive correlations, respectively. The diagonal of 1s has been removed. The genes are sorted using hierarchical clustering to reveal block structure at E11.5. Block structure disappears when genes are sorted the same way at later developmental stages, indicating that the correlational structure present at E11.5 is no longer present.

**Figure 2:**
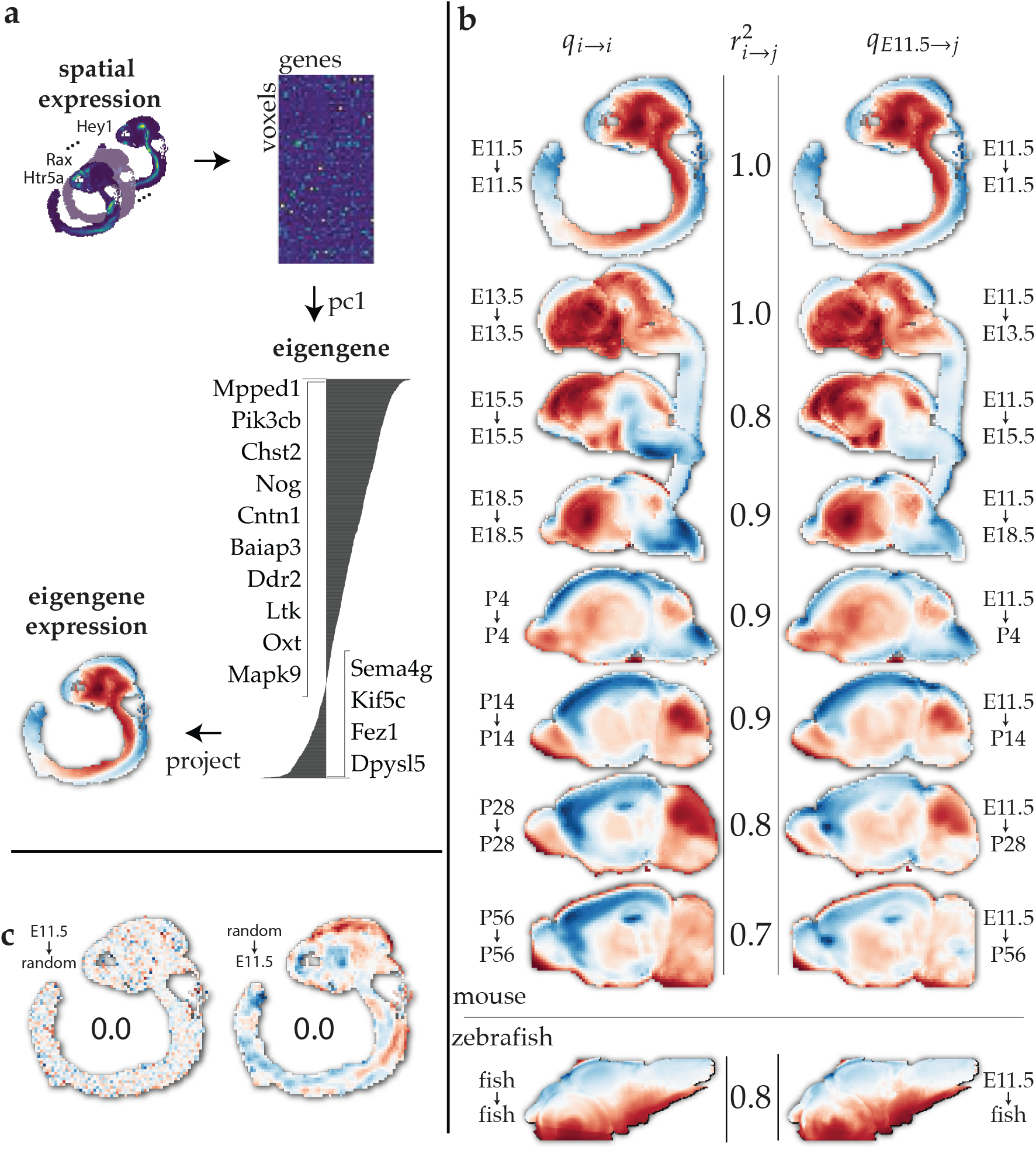
Spatio-temporal expression of the principal eigengene is stable across development and evolution. (**a**) Construction of principal eigengene for E11.5. The spatial expression pattern for all measured genes (upper left) is converted into the gene expression matrix *X*_*t*_ (genes × voxels). Following singular value decomposition of this matrix, the principal eigengene is extracted, and the spatial expression pattern is color-coded, with voxels in which the loading is positive shown in blue and those in which it is negative shown in red (lower left). (**b**) The principal eigengene computed at each developmental stage has the same expression pattern when applied at later stages. The eigengene computed at a given stage partitions the brain into two contiguous global regions, roughly along the dorso-ventral axis (left column). The E11.5 eigengene induces almost the same partitioning (*r*^2^ *>* 0.7) when applied to later stages or even onto the zebrafish (right column). This result holds for any combination of source eigengene and target stage (see Fig. 3a). (**c**) The eigengene measured at E11.5 does not reveal any structured expression pattern on randomized data (left), nor does an eigengene with randomly drawn loadings (right). In both cases, there is no correlation with the original pattern (panel c, top row), *r*^2^ = 0.

### Eigengene expression patterns are stable across development and evolution

We asked whether some characteristics of the pattern of gene expression, not evident at the level of single genes, might show greater stability across development. The data from a particular developmental stage *t* can be summarized as the gene expression matrix *X*_*t*_, an *n*×*m* matrix with *n* voxels and *m* genes (Fig 2a). In this matrix, each row represents a voxel,

which corresponds to a specific location in the tissue, and each column represents a gene, with the entries indicating the expression levels of that gene in the respective voxels. Note that although the matrix *X*_*t*_ contains all of the information about gene expression at a given developmental stage, it does not retain any information about the spatial organization of voxels.

The basis of our analysis is the eigengene, which is defined as the principal component of the gene expression matrix *X*_*t*_ (Fig 2b). In principal components analysis (PCA), the gene expression matrix *X* is decomposed into

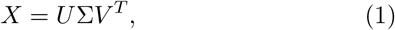

where *V* is an *m* × *m* matrix of loadings in which each row corresponds to a gene and each column constitutes a principal component; Σ is an *n* × *m* diagonal matrix containing how much variance is explained by each component; and *U* is an *n*×*n* matrix whose entries contain the coefficients that, when scaled by Σ, reconstruct *X* in its new basis *V*. We focus on the first principal component, PC1, at each stage,

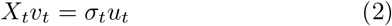

where *t* indicates the developmental stage (E11.5, E13.5, …), *v*_*t*_ is the first column of *V*_*t*_, also called PC1 and which we will refer to as the *principal eigengene* (Fig. 2b), and *Σ*_*t*_*u*_*t*_is the first column in *u*_*i*_ scaled by the first entry along the diagonal of Σ_*t*_. Consistent with the fact that *X*_*t*_ does not contain information about the spatial organization of voxels, the principal eigengene is also independent of the physical locations of the voxels: The loading profiles are identical if the voxels’ positions are shuffled before computing the principal component.

The principal eigengene can be thought of as a virtual genetic probe which detects the degree to which that pattern of genes is expressed in corresponding relative ratios, e.g., gene A is expressed twice as strongly as gene B and inversely correlated with gene C, in any given cell. The probe reports strong blue when there is an exact match between the principal eigengene and the genes expressed in a cell, and lighter blue for less perfect matches. If there is no match the probe is white. In cells in which the match is the exact opposite, the probe is red.

Fig 2b (*left*) shows the spatial loadings of the nine principal eigengenes—eight drawn from across mouse developmental stages and one for zebrafish–over the course of development. Casual inspection reveals strong spatial structure in the loadings: the brain at each stage is divided into about two to four contiguous areas, in marked contrast to the expression of individual genes, which tends to be concentrated in multiple hot-spots. Note that, because the principal eigengene is computed in expression space and not physical space, there is no *a priori* reason that individual voxels should show any spatial organization at all. The dominant feature of the spatial expression of the principal eigengene is the partitioning of the embryo globally, roughly along its dorso-ventral axis.

We next asked about the projection of the principal eigengene computed at one developmental state *t* to another developmental state *t*^′^. Just as the expression of the principal eigengene in each voxel can be found through the product *X*_*t*_*v*_*t*_, we define the expression of a principal eigengene *v*_*t*_ computed at one developmental stage and assessed at another stage *t*^′^ through the product

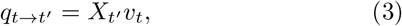

where *q*_*t*→*t*_′ is the expression of the eigengene at stage *t* onto the voxels of stage *t*^′^.

Strikingly, the expression pattern of the E11.5 eigengene applied to later developmental stages yields almost exactly the same dorso-ventral partitioning (Fig 2b, *right*). Moreover, the E11.5 principal eigengene is is not special: The projection between any pair of source and target stage yields similar spatial patterns (Fig. 3a), quantified as the squared correlation

**Figure 3:**
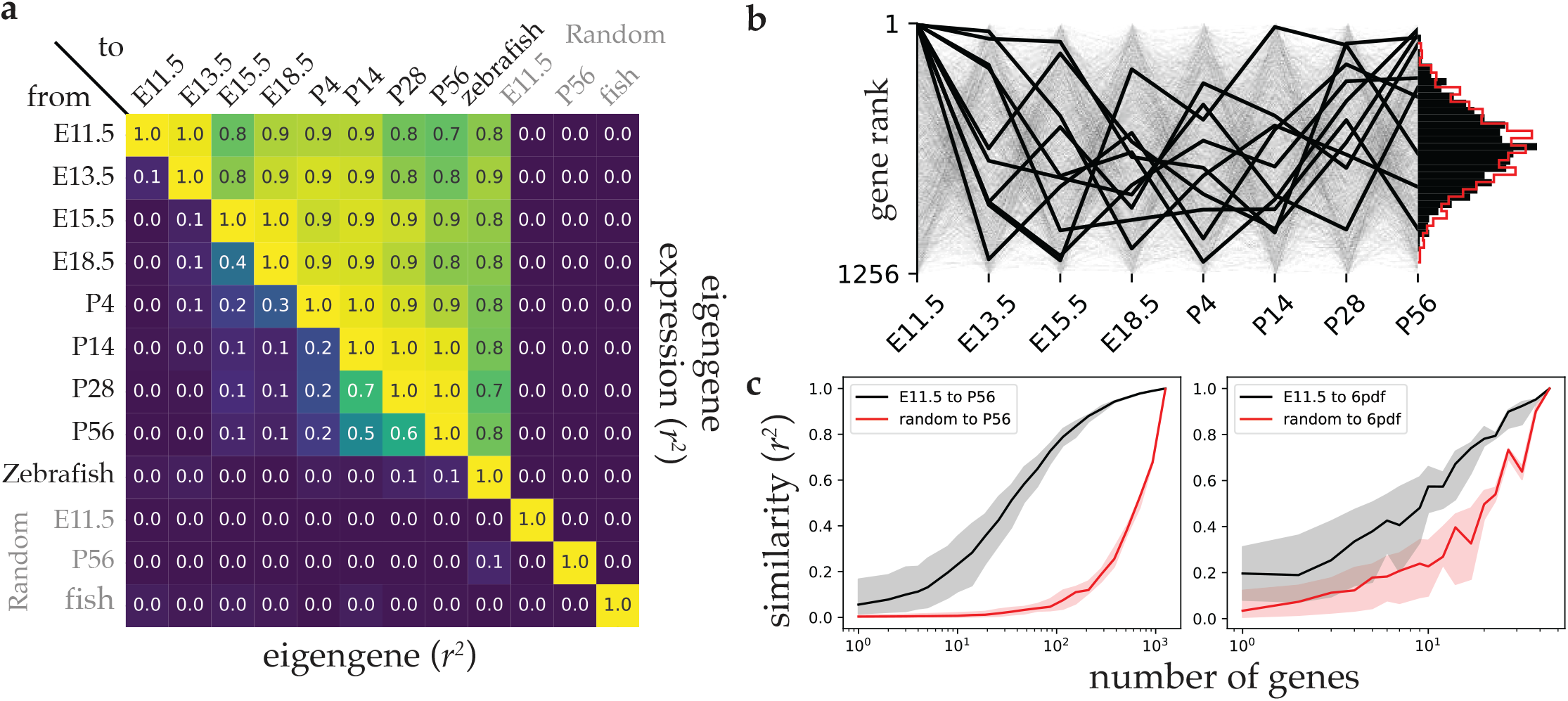
**The aggregate spatio-temporal expression of small random sets of genes correlates with the eigengene expression pattern a** The squared correlations of eigengene loadings across different stages (lower triangle) and their spatial expression (upper triangle). The matrix diagonal shows the correlation of each eigengene with itself, which are trivially 1.0 for both in terms of their loadings and their spatial expression, and the top row shows the projection of all stages onto the E11.5 eigengene (also shown in Fig. 2b). “Random” rows and columns represent randomly drawn eigengene loadings and expression voxels, respectively. The large values of the upper triangle indicate that spatial expression is stable whereas the low values in the lower diagonal indicate that the eigengene itself changes across development. **b** The rank of each gene in the eigengene loading vector is not consistent across developmental stages. Each line represents a gene, with bold lines indicating the top ten ranked genes at E11.5. The histogram of average ranks per gene matches the mean rank distribution for random ranks (shown in red). **c** Random subsets of genes recompute the eigengene at E11.5, demonstrating robustness. The *r*^2^ similarity of P56 and 6dpf zebrafish voxels projected onto this recomputed eigengene (black line) is compared with an eigengene with random loadings (red line). The subsets sampled at each size are non-overlapping, ensuring no gene is included in multiple sets of the same size.

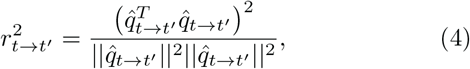

where 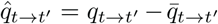 is the mean-subtracted version of *q*_*t*→*t*_′. To test whether this dorso-ventral partitioning somehow arises as the inevitable consequence of this projection operation, we projected onto random gene expression vectors instead of the principal eigengene as a control. However, these global patterns are not present when the analysis is applied to randomized expression data, nor when the voxels are projected onto a random axis rather than the eigengene (Fig 2d,3a). These results reveal that a signature of that the brain’s gene expression covariance is stable throughout the course of development.

Even more strikingly, the eigengene measured at mouse E11.5 not only preserves its spatial expression pattern across mouse development, but also across more than four hundred millions years of evolution since the mouse and zebrafish shared a common ancestor (Fig 2c). The projection from mouse to zebrafish was performed by selecting all homologous genes in zebrafish data from the genes available in the mouse data, without recomputing any loadings. Using this selection, the eigengene measured in E11.5 also elicits a dorso-ventral partition in the larval zebrafish brain. This result suggests that the specific spatial covariance pattern among genes is preserved across much of the vertebrate lineage and perhaps beyond.

The partitioning induced by the principal eigengene is similar to that caused by dorso-ventral patterning events, driven by genes such as Shh and BMP, early in development. Because our data begin at E11.5, considerably after these events, they cannot address the initial establishment of these patterns. However, the data suggest that whatever mechanisms establish the dorso-ventral gradient initially, the eigengene expression pattern is maintained through inheritance in absence of the original inducers, indicating a robust and conserved framework for spatial organization throughout development.

The preservation of the spatial covariance pattern could be explained if the eigengene’s loading vector itself remained constant throughout development. However, as shown above (Fig. 2a), the expression of typical single genes tend to fluctuate strongly over the course of development. Indeed, we observe that the loading vectors measured at different time points are largely uncorrelated (Fig. 3a,b). Despite their lack of correlation, their spatial expressions are very similar (*r*^2^ *>* 0.7). This observation presents an apparent paradox: How can the axis of covariance change, while its spatial projection remains constant?

A mathematical fact resolves this paradox: The expected correlation of two random vectors becomes very low for high dimensions. The expected *r*^2^ for two random vectors of 1000 genes is on the order of 1*/*1000^2^ = 10^−6^. Thus, the observed correlations, despite being small in single-variate terms, are not negligible in multi-variate terms. However, despite there being no mathematical contradiction, the correlations do indicate that loadings of the eigengenes rotate much more over time than their corresponding spatial expression patterns.

We next asked whether a small set of genes, such as those known to be involved in dorso-ventral patterning, are responsible for the emergence of this global spatial pattern. The natural metric of importance is the rank of a gene’s loading coefficient in the eigengene. However, we find that these ranks are essentially random over the measured eigengenes (Fig. 3b). Moreover, the conservation of the mouse projection to zebrafish using only a small subset of homologous genes could be explained if the spatial patterns are not contained in the covariance of just a few specific genes, but are instead spread across many subsets of genes.

To test this possibility, we measured the spatial patterns revealed by random subsets of genes of various sizes. We find that random subsets generate the same eigengene expression pattern, regardless of the selected genes, even when the subsets have no genes in common (Fig. 3c). Seemingly, the only requirement is that the subset be sufficiently large. This result indicates the spatial pattern information is broadly distributed across genes.

### The lineal model explains eigengene expression patterns

Sydney Brenner contrasted two models by which cells assume their identity during development, which he termed the *American Plan* and the *European Plan*. The European Plan, named after the notion that Europeans inherit their identity from their lineage, emphasizes the role of cell lineage in determining cell identity and thus gene expression patterns. In this model, gene expression patterns of daughter cells are similar to that of their parents and thus largely determined by each cell’s developmental history. According to the European Plan, spatial correlations in gene expression arise mainly from inheritance through cell division, and the fact that most cells don’t migrate very far, rather than from local cell-cell interactions (Kerstjens et al., 2022). By contrast, the American Plan, named after the idea that Americans derive their identity from the neighborhood in which they live, posits that local interactions between cells, mediated by signaling molecules and gradients, determine gene expression patterns. In this model, a cell’s fate and gene expression are determined by its local environment and the signals it receives from neighboring cells. The American Plan emphasizes the importance of cell-cell communication and the role of morphogens in establishing spatial patterns of gene expression. In what follows we will refer the European plan as the *lineal model* and the American plan as the *neighborhood model*.

We have shown so far that the expression of the principal eigengene partitions the brain along the dorso-ventral axis throughout development and into adulthood, and that this pattern of gene expression is preserved over more than four hundred million years of evolution. How does this pervasive spatial pattern arise? One possibility is that the spatial correlations arise from the neighborhood model, according to which communication among nearby cells drives them into a pattern of similar gene expression. Alternatively, the persistence of the principal eigengene across development could arise from the lineal model, as a direct consequence of constraints on mitotic division. To compare these two possibilities, we formulated simplified models of the lineal and neighborhood models.

In the lineal model (Fig 4a–c), a cell’s expression is the expression of its parent plus a differential expression vector that is drawn from a normal distribution. For a parent cell *c*_*i*_ with daughter cells *c*_*i*0_ and *c*_*i*1_, the expression of the daughters are

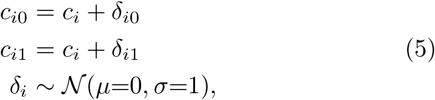

where *δ*_*i*_ is a vector of differential expression that is normally distributed with zero mean *μ* and unit standard deviation *Σ*. Note that the *δ*_*i*_’s are not random variables. Rather, they model the complex yet deterministic changes in gene expression between parent cells and their daughters. These changes are assumed to be distributed normally, but not drawn randomly. To model the spatial layout of cells we use an H-tree fractal, in which spatially proximal cells tend to be closely related in lineage (Espigulé, 2013; Mandelbrot, 1983).

**Figure 4:**
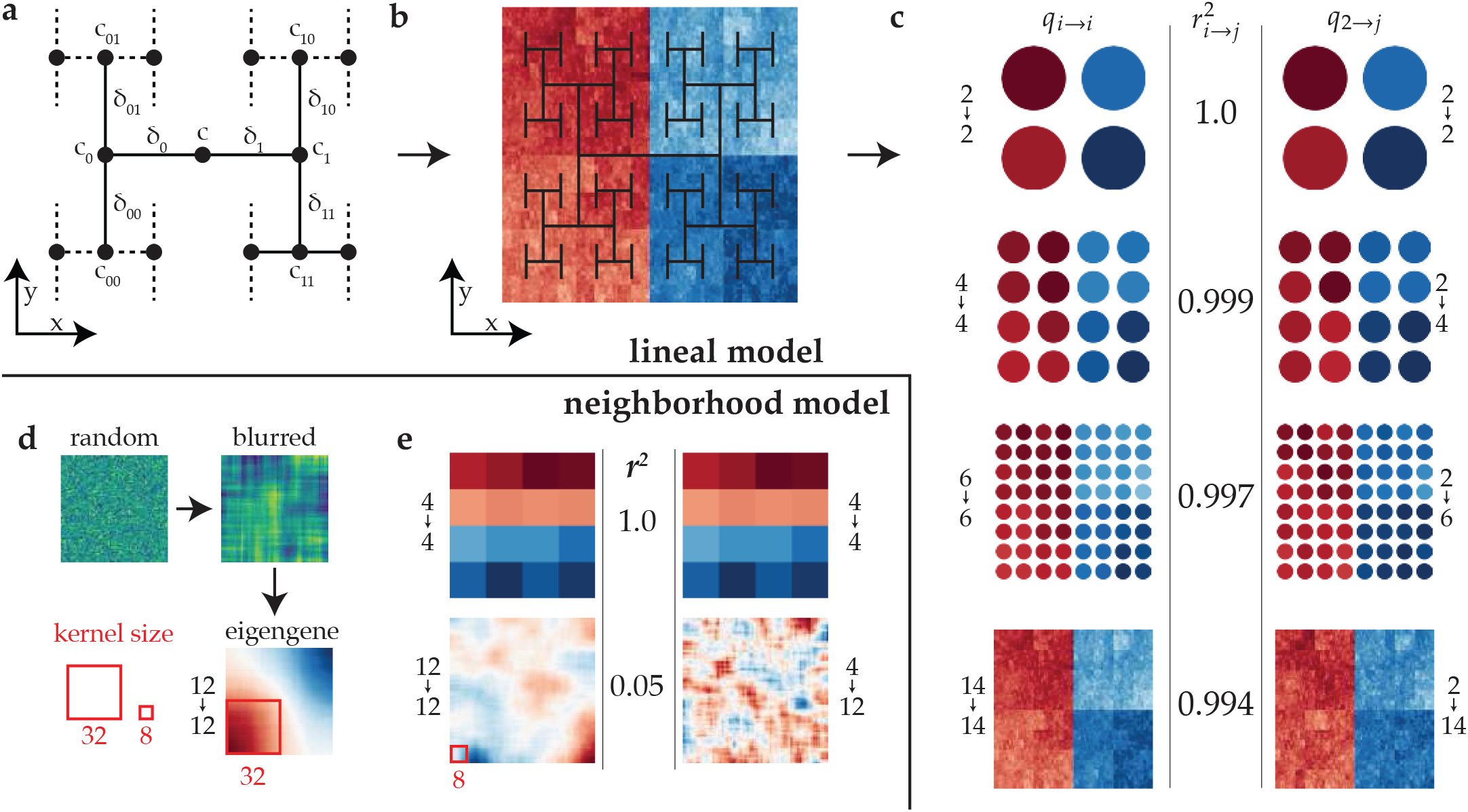
Simple lineal constraints can explain global eigengene patterns, whereas cell-cell interactions alone do not. (**a**) The lineal model is captured in a lineal model of gene expression. The gene expression *c*_*i*_ of cell *i* is modeled as a normally distributed deviation *δ*_*i*_ ∼ 𝒩 (*μ*=0, *Σ*=1) from its mitotic parent (See Eq. 5). The lineage tree is laid out in the spatial *x, y*-plane according to an H-tree fractal. (**b**) The leaves of a simulated lineage tree of 14 generations (2^14^ = 16384 cells), with each cell expressing 1000 genes. The leaf cells of this tree are collected, and the eigengene is measured across all leaves. The sign of the eigengene expression (red, negative; blue, positive) partitions the leaves into two spatially contiguous groups that correspond to the respective progenies of *c*_0_ and *c*_1_. The split in mitotic lineage thus implies a global partition both in gene expression space, as measured by the eigengene, and in physical space. (**c**) Multiple stages of development were generated by measuring the leaves of the simulated lineage after 2, 4, 6, and 14 rounds of division, analogous to the stages of mouse development. Then, the analysis of Fig. 2c was repeated for the simulated expression data. As in the experimental data, the principal eigengene measured at an early stage patterns the simulated tissue at the late stage, even after 14 generations of divisions. (**d**) The neighborhood model is captured in a local cell-cell interaction model of gene expression. An initially random grid is generated with 64 × 64 = 4096 cells that each express 1000 genes. The grid is then spatially blurred to induce spatial correlations. The principal eigengene across these simulated expression data only globally pattern the tissue when the range of the blur is large enough to break the assumption of locality, here with a width and height of 32 cells, i.e., half the size of the space. (**e**) However, when the range of the blur does not scale with the size of the tissue—this panel shows a range of 8—the eigengene expression does not form a global pattern, but rather a collection of blotches comparable to the random eigengene control in Fig. 2c). Furthermore, the local interaction model does not preserve the eigengene from early development (top row) to late development (bottom row): Comparison of the left and right column of the bottom row shows that when the early eigengene is projected to late expression data (4 → 12), the spatial (12 → 12) is not preserved (*r*^2^ ≈ 0.05).

In the neighborhood model (Fig 4d–e), the expression of each cell is drawn from a normal distribution with *μ* = 0 and *Σ* = 1, without regard for lineage. Then, the space of voxels is ‘blurred’ by performing a moving average window over spatial neighbors, causing neighboring voxels to be correlated. This blurring model intends to capture spatial correlations induced by local interactions among cells.

In simulations of both models, we find that the principal eigengene partitions the developing brain into regions (Fig. 4b,d). However, there are some important differences between the two models. First, in the lineal model global partitioning arises at every simulated developmental stage (Fig. 4c). In the neighborhood model, by contrast, the range of interaction is determined by the diffusion-mediated “blurring” (Fig. 4d,e), which causes nearby cells to be similar. So, to preserve a global pattern, the range of interaction must scale as the tissue grows. For large tissues, the interaction range may be larger than is physiologically plausible through simple diffused morphogens (Goodhill, 2016). Thus, the neighborhood model must invoke additional mechanisms for different sized brains (over development and across species) to explain the global partitioning, whereas the global partitioning is a natural prediction of the lineal model.

Second, the neighborhood model does not trivially produce a consistent pattern over time, especially as the tissue grows in size. This is because the characteristic frequency of the spatial pattern is a function of the interaction radius, and so for the frequency to decrease, the radius must increase as the tissue grows (Fig. 4d). By contrast, by virtue of the hierarchical nature of growth across generations in the lineal model, the model is intrinsically scale-free, and explains the temporal persistence and scaling (Fig. 4c).

Beyond scalability, the lineal model predicts that the hierarchical nature of the growth process is installed in the tissue as a hierarchy of nested spatial regions, each with a gene expression signature measurable as the eigengene that dominates the variance in that region. To assess whether the experimental expression data shows evidence of a hierarchical growth process, we performed an iterative hierarchical decomposition, similar to a hierarchical clustering. The method used here is adapted from (Kerstjens et al., 2022). First, the eigengene across all voxels is calculated, as in Fig. 2b. Then, the voxels are divided along the eigengene expression into two halves—namely those with positive and negative coefficients; and the main axis of variance is calculated within each of these halves. We repeat this process recursively until the tissue has been divided down into single voxels. This recursive process results in a tree of voxel subsets: The root of the hierarchy contains all voxels, and each leaf exactly one voxel.

We find that this hierarchy indeed exists in both the mouse and zebrafish brain, as well as in our lineal model (Fig. 5a). By contrast, a hierarchy only exists in the interaction model if the interactions occur over long large distances. As such, the hierarchy produced by the interaction model is not scale-free (Fig. 5a). We have quantified the hierarchy by the contiguity of the sub-regions (Fig. 5b). Each node of the hierarchy corresponds to a collection of voxels that are positioned in 3D physical space. These voxels may either be all contiguous, i.e., there exists a path between each pair of voxels passing only through immediately adjacent voxels, or be a collection non-contiguous voxels. We measure the contiguity as the size of the largest connected component divided by the total number of voxels in the region. This yields a number between 0 and 1, where 0 indicates there are many individual voxels that are all disconnected, and 1 indicates all voxels are contiguous (Fig. 5c). We find that our lineal model and both the zebrafish and mouse data create hierarchies of contiguous regions, whereas the interaction model does not (Fig.5b).

**Figure 5:**
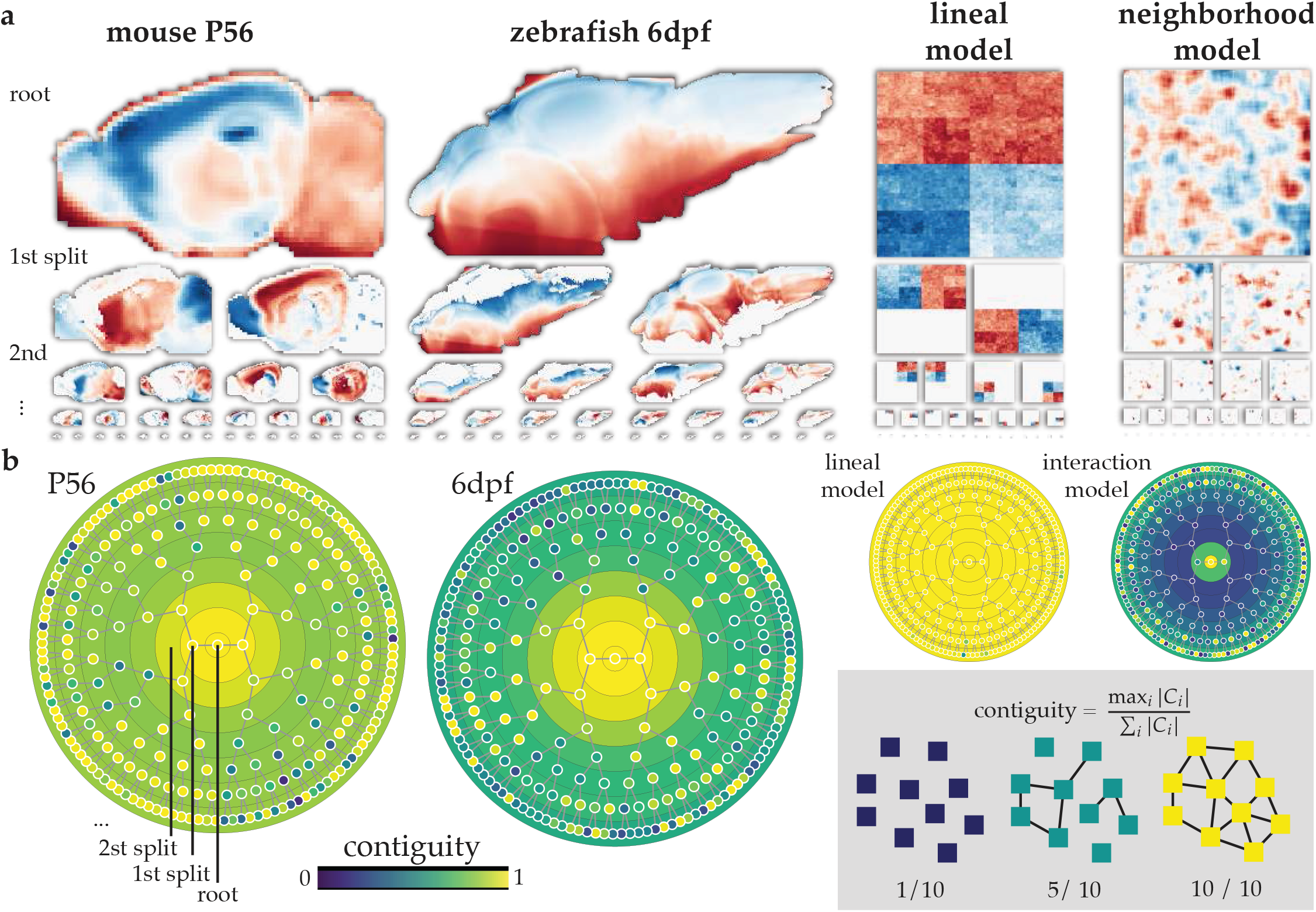
The developing brain is recursively partitioned into a hierarchy of eigengenes. (**a**) Eigengene expression splits the mouse and zebrafish brains in two regions, red and blue. We then measure the two eigengenes across only the voxels in one region, i.e., either only the red or only the blue voxels, and measure the eigengene expression within the regions. This yields a red-blue split within each region. We apply this procedure recursively to produce a hierarchy of nested regions. The figure shows only the first few generations, but the hierarchy continues until each region consists of only a single voxel. (**b**) The contiguity of each region is quantified. We define contiguity as the number of voxels in the largest connected component divided by the total number of voxels. The contiguity is 1.0 if all voxels are connected. If none of the voxels are connected, then the contiguity asymptotes to 0 as the number of voxels goes to infinity. Each node in the tree corresponds to a region. The center node is the root of the hierarchy, which trivially has a contiguity of 1 because it contains all the voxels. The leaves of the hierarchy (not shown) also have a contiguity of 1 because they all contain a single voxel. Each concentric circle corresponds to a tier in the hierarchy. The color of the tier is the average contiguity of the regions within the tier. The lineal model generates a perfect contiguity. The blurred control has poor contiguity. Zebrafish and Mouse brains have high contiguity in select branches, and a higher average contiguity than the blurred control case.

### Eigengene hierarchy enables efficient axon navigation

We have identified a global hierarchical feature of spatial gene expression that is conserved across mouse and zebrafish brains. Building on a previous mechanistic model of axonal navigation (Kerstjens et al., 2022), here we provide an intuition for how this hierarchy might be more than an epiphenomenon of lineal development, by providing the foundation for an efficient mechanism for wiring the brain.

The brain is wired as billions of neurons extend their axonal arbors to make trillions of connections. Each branch of the axonal arbor is guided by the growth cone at its tip, which measures subtle molecular gradients across the width of the cone. Molecular cues are not intrinsically attractive or repulsive. Rather, the cone’s molecular pathways decide which gradient to ascend or descend based on the cone’s intracellular state and extracellular context. To travel over longer distances, growth cones re-configure themselves—e.g., by exchanging its receptors—to tune into a different set of molecular gradients, or change the valence of the same cue. Axonal journeys are split into multiple legs by such re-configurations, which are triggered by extracellular cues and mediated by molecular pathways (Dorskind & Kolodkin, 2021; Goodhill, 2016; Stoeckli, 2018).

How might the mechanism that guides the individual axonal branches scale to wire a whole brain? The straight-forward strategy is to treat every axon individually (Fig. 6c). That is, each growth cone is guided to its target by its own stereotyped sequence of configurations. However, this strategy does not scale to large brains. For example, if a growth cone needed to navigate an axonal branch through ten regions, it would need to re-configure itself ten times. Each of these re-configurations requires a molecular pathway that is triggered by a cue in the growth cone’s environment, and effects a change in its internal state. A second growth cone navigating to a different target might need to re-configure itself with ten different configurations that follow slightly different rules and so employ slightly different molecular pathways, even if it is a branch of the same axon. As the number of growth cones and targets increases, the number of re-configurations that must be encoded grows to become prohibitively large. Even if receptors and ligands are re-used, the genome cannot explicitly encode all growth cone re-configurations: With *k* legs per growth cone, *m* targets per axon, and *n* neurons, the total number of re-configurations would be the product of these three numbers: *nmk*. For a human brain with *n* ≈ 10^10^ neurons, *m* ≈ 10^4^ targets per neuron, and *k* ≈ 10 legs per journey, the number of growth cone re-configurations is at least of the order ∼10^15^. With only ∼10^9^ base pairs, the human genome cannot explicitly encode triggers and pathways for all growth cone reconfigurations. So, we must consider more efficient strategies to explain how large numbers of axons are informed of their individual routes.

**Figure 6:**
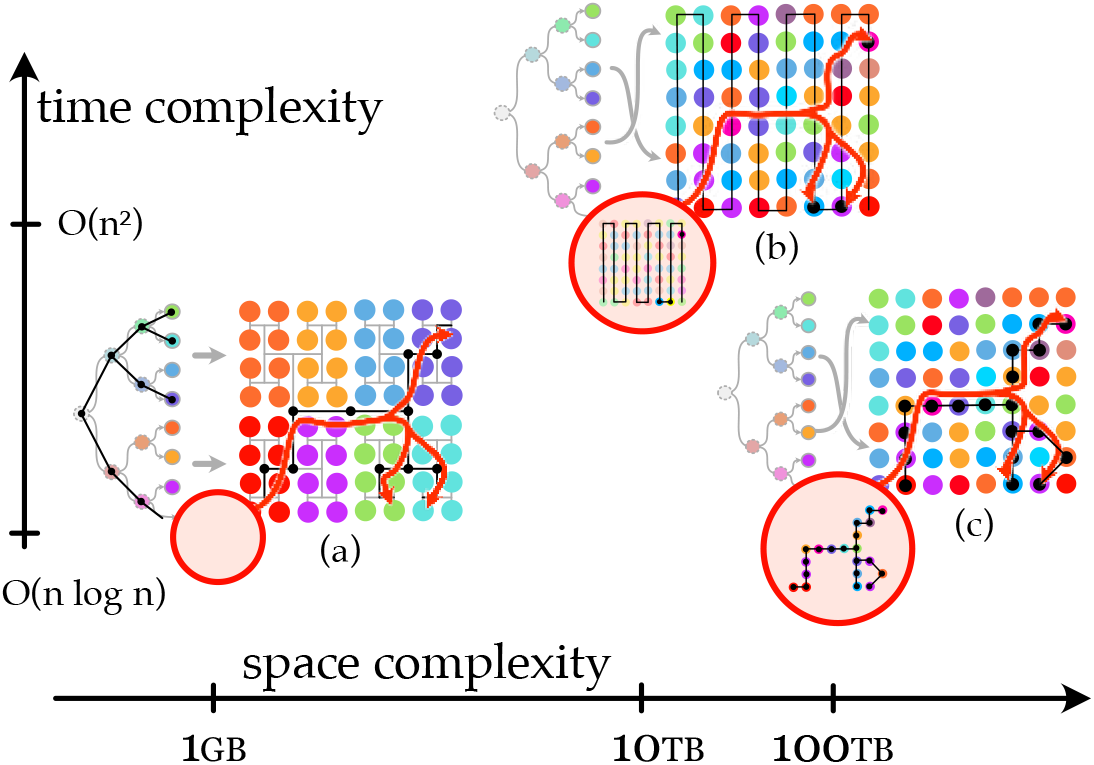
Three potential strategies for axon navigation with different time and space complexities. Strategies (b) and (c) navigate through a field of cells (circles) with various gene expression signatures (colors) without any spatial structure among them. Strategy (c) treats all axons independently, and thus stores the routes to all targets in each axon. The space complexity of this strategy is *O*(*nmk*), where *n* is the number neurons, *m* the average number of targets per neuron, and *k* ≪ *n* the average number of legs in a journey from a source neuron to a target. The time complexity is also *O*(*nmk*), as every axon branch finds its targets in *k* ≪ *n* steps. In strategy (b) the routes are not stored, and so the space complexity is reduced by a factor *k* to *O*(*nm*). Instead, the axon finds its targets by systematically visiting every cell in the tissue. However, each neuron needs to visit every other neuron, which implies a time complexity of *O*(*n*^2^). Strategies (b) and (c) is only tractable at small scales. Strategy (a) assumes the tissue is organized as a lineage-induced hierarchy, whose identification and mechanism of generation are the main topics of this paper. Exploiting this organization allows an axon to find its target in *O*(log *n*) steps by effectively tracing the lineage, bringing the whole algorithm to a time complexity of *O*(*nm* log *n*); and, it does this with no additional data over what has been already described to differentiate cells to their specific fates, so the space complexity amortizes to *O*(1). The details of this navigation algorithm are published in (Kerstjens et al., 2022).

Computer science studies the efficiency of strategies, and uses the notion of *algorithmic complexity* to speak about such questions. The *space complexity* of an algorithm is the the relation between the size of the problem at hand, and amount of data the algorithm uses to solve it. For example, the space complexity of wiring the brain by explicitly encoding all growth cone re-configurations is *O*(*nmk*), as described above. (The big-*O* denotes that *nmk* is not the exact amount of data requires, but that the real amount scales asymptotically as *nmk*.) Analogous to space complexity, there is also time complexity, which captures how many steps are needed to execute the strategy. The time complexity for the explicit encoding strategy would also be *O*(*nmk*), because *n* neurons need to take *k* navigation steps for each of *m* axonal branches. An algorithm is *tractable* if both its time and space complexity are sufficiently small. In our case, the global wiring algorithm must fit in the genome and execute within the period of gestation.

The essential issue with the explicit encoding strategy (Fig. 6c) is not that each axon needs to store re-configurations for the *k* legs of its journey. One could imagine a strategy whereby the axon finds its target by systematically searching the whole brain, completely eliminating the need to encode the source-target trajectory (Fig. 6b). This unrealistic strategy would take a prohibitive amount of time to execute, as each of *n* neurons would need to visit every other neuron to check whether it is one of its targets, implying a total time complexity of *O*(*n*^2^). However, even this time-intensive scheme would only reduce the space complexity with a factor *k*, because each of *n* axons would still need to encode a list of all its *m* targets, yielding a space complexity of *O*(*nm*). That is, the amount of encoded information still scales with the total number of connections *nm*, which is of the order 10^14^—still 5 orders of magnitude larger than the information storage capacity of the genome.

It follows that growth cone re-configurations must derive from a simpler strategy whose time and space complexities are not proportional to the number of connections in the brain. In previous work (Kerstjens et al., 2022) we proposed an axonal navigation strategy whereby growth cones do not reconfigure using their own locally stored regulatory information, but rather co-opt the information already in place through cell differentiation (Fig. 6a). The intuition behind this strategy is that the blueprint for building a brain region can also be used as a map to navigate it. If a growth cone can partly recapitulate the regulatory logic of differentiation, it could re-use it to generate growth cone configurations for navigation. The question of which regions are traversed then reduces to which differentiation programs the cone can successfully recapitulate. Because neither the axon’s targets nor their routes are explicitly encoded, this strategy incurs only the fixed information cost (space complexity *O*(1)) needed to convert differentiation logic to navigation logic. In other words, the information required to navigate axon using this scheme does not grow with the size of the brain, which could explain how the same strategy can be used to wire a ∼1g mouse brain and a ∼1kg human brain.

However, this efficient navigation scheme requires that the brain’s tissue be organized as the lineage tree, along which cells differentiate. In this paper we offer a mechanism whereby the lineage tree embeds its hierarchical structure onto the physical and epigenetic relations among cells (Figs. 4,5). In other work we have detailed how axons might use such a space for efficient navigation (Kerstjens et al., 2022).

The model presented here (Fig. 4) does not specify how differentiation is regulated, nor how it informs the growth cone. It merely demonstrates a mechanism whereby lineal constraints provide an organized space that supports at least one axon navigation strategy whose information cost does not grow proportionally to the number of connections in the brain^1^. Future models might capture the phenomenological constraints we assert here in a more mechanistic model of the gene regulation network of differentiation, and the molecular pathways that allow its re-use in axon navigation.

## Discussion

We have analyzed the spatial expression of genes across mouse brain development and in larval zebrafish. We find that, even though the expression of individual genes is highly dynamic, a simple function of gene expression—the principal eigengene—is remarkably conserved across space, ontogeny, and phylogeny. This eigengene can be recovered even when computed from only a small random subset of genes, indicating that it is not the direct result of a small number patterning genes. These features can be explained by a simple model of inheritance in which the transcriptomes of daughter cells tend to be similar to that of their parent, but not by a simple model of cell-cell interactions alone. We speculate that the global nature of the eigengene hierarchy might provide a conserved global strategy for imparting positional information that could be used for long-range developmental processes such as pattern formation and axon guidance.

### Eigengenes in mice and fish

We observe in both mouse and zebrafish that the measurement of expression covariance across any sufficiently large set of genes at a single developmental stage yield an eigengene that reveals a stable pattern of differential gene-expression across the entire brain (Fig. 2c). The pattern is hierarchical (Fig. 5), and so ranges in scale from single voxels to the whole brain. The pattern is stable over all available developmental observations (Fig. 2c,3a). Moreover, the expression covariance measured in mouse can be used to predict the spatial pattern observed in a fundamentally different species, the zebrafish (Fig. 2c).

This result is surprising for a number of reasons: Firstly, our analyses do not use explicit spatial information: The eigengene is measured across the full collection of data voxels without regard to their *x, y, z*-location. Therefore, whatever spatial information is present in the data, must be implicit in the gene expression itself and not entailed by our analysis. This is confirmed by our computational controls (Fig.2d).

Secondly, the hierarchy of eigengenes is temporally static and spatially global, despite the dynamic and local nature of gene expression during development, and the many streams of cell migration. Establishing such global structure is not trivial, as biological development lacks an external constructor that can control development from the outside and provides a global reference frame. Instead, cells must negotiate global structure through their individual local behaviors.

One hypothesis as to how non-local structure might emerge is through the emission of molecular signals that diffuse over long distances. However, the effective range of a diffused signal is insufficient to establish global structure (Goodhill, 2016). Other mechanisms for establishing long range gradients exist, but require complex regulation to establish molecular relays and amplifiers. From this perspective, it is surprising to find a global organization of gene expression that spans all measured genes and stages of development.

Thirdly, the eigengene is agnostic to the specific genes that are measured (Fig. 3). The default hypothesis for the emergence of a spatial pattern across a developing tissue is for there to be a specific molecular pathway that directly regulates molecular gradient, which in turn effects changes in downstream targets. Although possible, it is not trivial that gene expression across all measured genes should align with some inducing gradient signal. However, we observe that the eigengene pattern is present across virtually any set of genes in the analyzed data. This implies that either the majority of genes align with a global morphogenic signal, or that some other mechanism, not relying on morphogenic signals alone, is responsible for the eigengene gradient. Note that this does not imply that the eigengene pattern is not subject to genetic regulation or morphogenic action. Rather, morphogenic action may leverage, consolidate, and/or shape the eigengene pattern that is induced through other mechanisms, such as the lineal model we propose here.

Fourthly, the eigengene expression pattern is preserved across mouse and zebrafish. It is known that the genes responsible for establishing the main body axes are highly conserved. For the dorsal-ventral patterning of the neural tube, for example, Shh and BMP are known players across species that have a notochord. However, we have not specifically selected Shh, BMP, or any of their known downstream targets; nor have we selected a developmental stage when these genes are known to be active. Yet the eigengene expression is conserved across an arbitrary selection of genes; at an arbitrarily selected developmental stage.

### Lineage induces an eigengene hierarchy

We considered two hypotheses for how global eigengene patterns may emerge. It is clear that nearby cells have similar expression, and that this can cause a covariance gradient. However, this observation alone does not answer why and how neighboring cells become similar. One could propose various mechanisms by which cells could negotiate similar cell expression through local interactions (Gierer & Meinhardt, 1972; Turing, 1952). Indeed, we have offered a phenomenological version of a neighborhood model (Fig. 4).

An alternative hypothesis, which we propose here, is that the eigengene variance axis is a direct result of constraints on cell division. Although this hypothesis also implies that nearby cells will have similar expression, our hypothesis is stronger as it predicts that cells of the same lineage retain a similar gene expression, even after many rounds of division. As such, it has a temporal aspect in addition to a spatial aspect.

There are several differences between the expression patterns that emerge from the two models. Firstly, the pattern induced by the lineal model is scale-free. That is, the tissue retains the eigengene expression pattern induced at an early developmental stage as it grows to an arbitrary size. That is, both the pattern’s size and characteristic space constant increase as the tissue grows (Fig. 4c). On the other hand, the local neighborhood model needs to adapt its action radius, i.e., the kernel size of the moving average window, as the tissue grows (Fig. 4d). This implies that cells would need to signal over much longer distances to form long-range similarities. Although conceivable, such signalling requires the adaptive regulation of the signal radius, and signal relays that overcome the limited distance over which morphogens diffuse. Such relay signalling may be implemented for specific genes, but it is harder to imagine how it would affect expression covariance across many genes (Stapornwongkul & Vincent, 2021).

Secondly, the lineal model explains how the eigengene measured at E11.5 can persist throughout the development of the organism (Fig. 4c). As the lineage tree grows, the original variance induced at an early stage is inherited by the progeny. Although each individual cell gradually loses similarity with its distant ancestor, the ancestral expression profile becomes shared among an increasing number of progeny. This balance retains the early variance between mitotic siblings as the principal component among their respective progenies. On the other hand, the local neighborhood model has no inherent mechanism that could retain the eigengene over time. More complex neighborhood interaction models that support specific regulation may be able to maintain an eigengene over time. However, the temporal maintenance of the eigengene would need to be explicitly regulated, whereas in the lineal model it is inherent.

Thirdly, the lineal model predicts that, beyond the global eigengene, each sub-region within the global eigengene (blue versus red in Fig. 2) has their own eigengene pattern that globally patterns the sub-region. This prediction applies recursively to generate a hierarchy of regions (Kerstjens et al., 2022). By measuring the first principal component recursively in this fashion, we uncovered a consistent hierarchy in both the mouse and zebrafish brains (Fig. 5). The model predicts that this hierarchy should be shaped as the lineage tree of cell divisions. However, testing whether the hierarchy follows true mitotic relations, e.g., through lineage tracing, is beyond the scope of the present paper. Interestingly, an eigengene hierarchy is also predicted by the neighborhood model (not shown) (Shinn, 2023). However, this hierarchy would not be temporally consistent, and require interactions at the scale of the whole organism.

Of course, the lineal and neighborhood models are not mutually exclusive. And, to some extent, a minor form of local interaction is already captured by the lineal model, namely the interaction between mitotic siblings that causes their differential expression. Future models should incorporate specific regulation, and aspects of both inheritance of expression state through the lineage and communication through interactions with spatial neighbors. Although these interactions were not necessary to explain the qualitative emergence of eigengene patterns, we expect they will be necessary to explain the specific geometry of eigengene patterns, which will depend on specific ‘traditional’ gene regulation.

### Global pattern formation

Our results raise the following view of global pattern formation:

During early development, all cells are close to one another, and each cell communicates with every other through short-range molecular signals. These cells behave as one tightly coupled dynamical system of intra- and inter-cellular dynamics, and collectively negotiate their respective gene expression. These dynamics induce a stereotyped spatiotemporal distribution of gene expression states across cells.

As the tissue continues to grow, cells will leave one another’s signalling range, so that the initial unified region splits into two adjacent regions with relatively independent dynamics. However, as cells inherit their expression state from their mitotic parent, the original spatial distribution of expression states, observed as the principal eigengene across the grand region, is preserved across the tissue as a whole. The subregions each establish their own distribution of gene expression states on top of the inherited distribution, and thereby layer an additional principal eigengene onto their domain of the global tissue (Fig. 4). This process is recursively repeated to establish a hierarchy of spatio-temporal eigengenes. Each induced eigengene acts as an invariance, as it is preserved throughout the rest of development, largely unaffected by subsequent changes in tissue shape, size, and gene expression.

Although an hierarchical view of embryonic pattern formation has been proposed in terms of hierarchical regulatory networks that cause hierarchies of progenitor fields (Davidson & Erwin, 2009; Davidson & Peter, 2015), we instead propose that hierarchical patterning is not the result of specific regulation, but in fact largely unavoidable in a tissue of dividing cells (Kerstjens et al., 2022). The surprising prediction from this counter-regulatory view is that the patterns should not be embedded in a small collection of dedicated patterning genes, but rather in the co-variance across essentially all differentially expressed genes. This prediction is confirmed in the specific spatial patterns of the eigengenes we have measured here.

An important open question is whether the eigengene has a mechanistic function in development, or whether it is an epiphenomenon of gene expression. The spatial patterning of the developing organism is a well-studied phenomenon. However, most, if not all, instances of such patterns (Briscoe & Small, 2015; Goodhill & Xu, 2005; Hubert & Wellik, 2023; Jaeger, 2011) are induced by a hand-full of specific morphogens and cover relatively small tissues. How do these morphogenic models of patterning (Gierer & Meinhardt, 1972; Turing, 1952) relate to the observation of a static expression hierarchy across eigengenes? And, could the hierarchy of eigengenes provide a global system for imparting positional information (Wolpert, 1969) to individual cells?

A global hierarchy of eigengenes is compatible with morphogens. One possible relation is that individual morphogens are representative factors of eigengenes. Such a relation would cast morphogens as the molecular mechanism for how positional information is imparted by the eigengene to individual cells. However, more complex relations, where single morphogens represent multiple eigengenes, are also conceivable. The exact relation between morphogens and eigengenes is an interesting avenue for future investigation.

The currently prevailing view is that there is no global system for imparting positional information. Instead, local coordinate spaces are separate, and long range developmental processes, such as long range navigation of axons, ‘stitch’ together a sequence of local spaces to form a global one. However, here we raise the possibility that there exists a global hierarchical coordinate space that the local spaces are sections of. Such a global structure would be especially useful in efficiently coordinating global developmental processes, such as long range migration of cells and axons (Fig. 6).

The theory of pattern formation hypothesized here raises the exciting possibility that the complex spatio-temporal dynamics among individual genes may be understood parsimoniously at the statistical level of their principal eigengenes that are invariant across ontogenesis and phylogenesis; analogous to how the complex interactions among many individual particles are understood parsimoniously at the statistical level of pressure, volume, and temperature.

## Methods

### Experimental data

The mouse data was collected by the Allen Institute(Thompson et al., 2014). From their data we have selected only the genes that have valid measurements across at least 90% of the voxels compiled across all time points, and only the voxels that have valid measurements across at least 90% of the remaining genes. This resulted in a set of 1256 genes.

Remaining missing values are filled with the mean expression value across the gene. This goes against the recommendation of the Allen Institute, which is to interpolate the value from the surrounding voxels. However, we decided to avoid risking any artificial induction of local similarities. Filling the value with the mean expression value does not introduce any additional variance.

Each gene’s expression is normalized to zero mean and unit variance within the voxels of their time point. This allows expression values to be both positive or negative, which we conceptually interpret as differential expression above and below a mean baseline.

The zebrafish data was collected by the Mapzebrain team (Shainer et al., 2023). The zebrafish data contains 290 genes. Of these genes, there is an overlap of 45 with mouse. The source data did not indicate valid versus invalid measurements, so the pre-processing step of excluding genes with high failure rates was not performed. Otherwise, all analyses are identical. Homologous genes were selected by choosing genes that have the same name in the mouse and zebrafish datasets.

### Eigengene

To calculate the eigengene, i.e., the global axis of variance, we use the ‘PCA’ routine of the ‘scikit-learn’ python package(Pedregosa et al., 2011). Although this routine calculates all principal components, we only consider the first.

Projecting a collection of voxels *X*, where the columns are genes and the rows the individual voxels, onto a component *ν*, is an ordinary matrix-vector multiplication *Xν* = *s* yielding the eigengene expression *s*.

### Hierarchical decomposition

The hierarchical decomposition was performed by recursively applying the eigengene analyses(Kerstjens et al., 2022). The first tier of the hierarchy is calculated as described in the previous section, and the voxels are projected back onto the eigengene to yield the eigengene expression *s*. This vector has one value per voxel. Then, all voxels are sorted based on whether their expression *s*_*i*_ (the ith voxel in expression vector *s*) is positive or negative. The negative voxels go into one bin, and the positive into another. The sign corresponds to the red (negative) and blue (positive) colors across our figures. The eigengene projection is then repeated for each bin independently. This process is then recursively applied untill only single voxels remain. Note that we do at no point calculate the second principal component, or remove any variance from the data. Each subsequent eigengene analysis is performed on the full expression data, without removing the variance of the previous eigengene analysis. The only difference is that the analysis is performed on subset of voxels.

### Connectedness metric

To measure the ‘connectedness’ of the data we employ the following measure. We first infer a geometric graph from the positions of the voxels in 3D space. For this we use a Delaunay tessellation (Delaunay, 1934; Virtanen et al., 2020) followed by applying the Gabriel criterion (Gabriel & Sokal, 1969). The result is a spatial graph where cells are connected to their immediate neighbors, while reducing the number of connections that cross open spaces. For example, in the case of a 2D images, as generated by our simulations, this procedure will make the expected square grid with adjacent pixels connected, but no connections across the diagonals.

A region of the hierarchical decomposition contains a collection of voxels. The spatial graph is established over all voxels. The voxels of a region make a sub-graph containing only the edges that have both their end-points in the regions. I.e., half-edges are not considered. The resulting spatial graph may be one large connected component. However, in general, the graph will consist of multiple collections of voxels that cannot be traversed between following only the remaining edges in the sub-graph. (We used the python package NetworkX (Hagberg et al., 2008) for its graph algorithms.)

Intuitively, if the region consists of one large connected component, the connectedness metric should be 1. If the region consists of independent singular voxels, the connectedness metric should go to 0. We chose a metric that depends on the number of voxels, so that a region with many independent voxels is closer to 0 than a region with few independent voxels. However, we wanted to avoid the case where the connectedness score of one giant connected component of many voxels would be dramatically reduced by a few single separate voxels.

To accommodate these features, we chose the following metric:

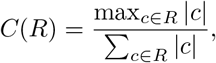

where *c* ∈ *R* are the connected components in region *R*, |*c*| is the number of voxels in that component.

### Lineal model

We describe the expression of a cell with index *i* as a vector

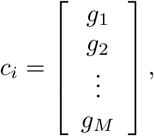

where each entry *g*_*j*_ is the expression of a hypothetical gene with index *j*. Although transcription measurements, such as RNA counts, are usually discrete and strictly positive, we allow real-valued expression for simplicity. These values can be thought of as deviations from a baseline average expression.

Genes are indexed 1 through *M*, where *M* is the total number of genes considered. Cells are indexed according to their position in the mitotic lineage. The initial cell at the root of the tree has no index. Its daughters are indexed 0 and 1. The daughters of cell 0 are indexed 00 and 01, and the daughters of cell 1 are indexed 10 and 11. This indexing scheme continues recursively, where in general the daughters of cell *i* are numbered *i*0 and *i*1.

We model the expression of a cell as a difference with its parent:

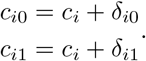

where *c*_*i*_ is the expression of a cell *i* that has two daughters *i*0 and *i*1, and *δ*_*i*0_ and *δ*_*i*1_ are the differential expressions between the parent and the daughters, respectively. This formulation is, in principle, without loss of generality, because each cell can be described in terms of a differential expression with its parent. This formulation does not preclude complex mechanisms of regulation or cell-cell signalling.

The generality is marginalized now we constrain the differential expression vectors *δ*_*i*_ are constrained to be normally distributed:

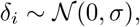

where *Σ* is the standard deviation of the distribution. This constraint implies that the daughters are one “step” away from their parent.

### Interaction model

The interaction model starts with a random matrix of size *n*×*n*×*M*, where *n*^2^ is the number of cells, and *M* the number of genes. The values in the matrix are normally distributed with a zero mean and unit standard deviation. This matrix is then blurred by applying a uniform convolution kernel of size *k* × *k* × 1. That is, each gene is independently blurred by replacing each value with *k* × *k* moving average window. The edges of the matrix, where the window would exceed the bounds of the sample, are padded by reflecting the values inside-out(Virtanen et al., 2020).

## Code & data availability

All code used for this work will be available upon final publication. All data is made available by the Allen Brain Institute (Thompson et al., 2014) and the Mapzebrain team (Shainer et al., 2023).

## Acknowledgements

The authors thank Arka Banerjee, Josh Dubnau, and Laura Andreae, for their useful discussion and feedback in writing our manuscript.

This work was supported by the Mathers Foundation (AZ) and the ETH Zurich Doc.Mobility Fellowship (SK).

## Footnotes

1 Computer scientists have long recognized that globally organizing a space of objects provides powerful algorithmic advantages. For example, we organize our English Dictionaries to be in alphabetical order so that we can find an arbitrary word in relatively few steps: If we want to find the word *epiphenomenon* we start by picking a word roughly in the middle of the dictionary, perhaps *mystery*. Now, we can reject every word that occurs after *mystery* because we know *epiphenomenon* will occur before *mystery*. Continuing, we can reject half the remaining words at each iteration, and find one word among a million in only 20 steps; this algorithm is called “bisection search”. In general, we will need at most *O*(log_2_ *n*) iterations to find any word among *n* sorted words. By contrast, if the dictionary were not sorted, there would be no other choice but to go through each word one by one until we found our target. This strategy would take at most *O*(*n*) iterations, and is the reason our dictionaries come pre-sorted.

## References

Briscoe, J., & Small, S. (2015). Morphogen rules: Design principles of gradient-mediated embryo patterning. Development, 142 (23), 3996–4009. 10.1242/dev.129452

Crick, F. (1970). Diffusion in embryogenesis [Publisher: Nature Publishing Group]. Nature, 225 (5231), 420–422. 10.1038/225420a0

Davidson, E. H., & Erwin, D. H. (2009). An integrated view of precambrian eumetazoan evolution. Cold Spring Harbor Symposia on Quantitative Biology, 74, 65–80. 10.1101/sqb.2009.74.042

Davidson, E. H., & Peter, I. S. (2015). Genomic control process: Development and evolution. Elsevier. 10.1016/C2012-0-02817-7

Delaunay, B. N. (1934). Sur la sphère vide. Bulletin de l’Académie des Sciences de l’URSS. VII. Série, 1934 (6), 793–800.

Dorskind, J. M., & Kolodkin, A. L. (2021). Revisiting and refining roles of neural guidance cues in circuit assembly. Current Opinion in Neurobiology, 66, 10–21. 10.1016/j.conb.2020.07.005

Espigulé, B. (2013). Generalized self-contacting symmetric fractal trees. Symmetry: Culture and Science, 24, 320–338. 10.26830/symmetry20131-4

Francis, K., & Palsson, B. O. (1997). Effective intercellular communication distances are determined by the relative time constants for cyto/chemokine secretion and diffusion. Proceedings of the National Academy of Sciences, 94 (23), 12258–12262. 10.1073/pnas.94.23.12258

Gabriel, K. R., & Sokal, R. R. (1969). A new statistical approach to geographic variation analysis. Systematic Zoology, 18 (3), 259. 10/fshrd8

Gierer, A., & Meinhardt, H. (1972). A theory of biological pattern formation. Kybernetik, 12 (1), 30–39. 10.1007/BF00289234

Goodhill, G. J. (2016). Can molecular gradients wire the brain? [Publisher: Elsevier]. Trends in Neurosciences, 39 (4), 202–211. 10.1016/j.tins.2016.01.009

Goodhill, G. J., & Xu, J. (2005). The development of retinotectal maps: A review of models based on molecular gradients [Publisher: Taylor & Francis eprint: 10.1080/09548980500254654]. Network: Computation in Neural Systems, 16 (1), 5–34. 10.1080/09548980500254654

Green, J. B. A., & Sharpe, J. (2015). Positional information and reaction-diffusion: Two big ideas in developmental biology combine. Development, 142 (7), 1203–1211. 10.1242/dev.114991

Hagberg, A. A., Schult, D. A., & Swart, P. J. (2008). Exploring network structure, dynamics, and function using NetworkX.

Hubert, K. A., & Wellik, D. M. (2023). Hox genes in development and beyond. Development (Cambridge, England), 150 (1), dev192476. 10.1242/dev.192476

Jaeger, J. (2011). The gap gene network. Cellular and Molecular Life Sciences, 68 (2), 243–274. 10.1007/s00018-010-0536-y

Kerstjens, S., Michel, G., & Douglas, R. J. (2022). Constructive connectomics: How neuronal axons get from here to there using gene-expression maps derived from their family trees (M. Kaiser, Ed.). PLOS Computational Biology, 18 (8), e1010382. 10.1371/journal.pcbi.1010382

Mandelbrot, B. B. (1983). The fractal geometry of nature (Updated and augmented) [OCLC: 36720923]. W.H. Freeman.

Pedregosa, F., Varoquaux, G., Gramfort, A., Michel, V., Thirion, B., Grisel, O., Blondel, M., Prettenhofer, P., Weiss, R., Dubourg, V., Vanderplas, J., Passos, A., Cournapeau, D., Brucher, M., Perrot, M., & Duchesnay, É. (2011). Scikit-learn: Machine learning in python. Journal of Machine Learning Research, 12 (85), 2825–2830. Retrieved November 17, 2022, from http://jmlr.org/papers/v12/pedregosa11a.html

Shainer, I., Kuehn, E., Laurell, E., Kassar, M. A., Mokayes, N., Sherman, S., Larsch, J., Kunst, M., & Baier, H. (2023). A single-cell resolution gene expression atlas of the larval zebrafish brain. SCIENCE ADVANCES.

Shinn, M. (2023). Phantom oscillations in principal component analysis [Publisher: Proceedings of the National Academy of Sciences]. Proceedings of the National Academy of Sciences, 120 (48), e2311420120. 10.1073/pnas.2311420120

Sperry, R. W. (1963). Chemoaffinity in the orderly growth of nerve fiber patterns and connections* [Publisher: Proceedings of the National Academy of Sciences]. Proceedings of the National Academy of Sciences, 50 (4), 703–710. 10.1073/pnas.50.4.703

Stapornwongkul, K. S., & Vincent, J.-P. (2021). Generation of extracellular morphogen gradients: The case for diffusion. Nature Reviews Genetics, 22 (6), 393–411. 10.1038/s41576-021-00342-y

Stoeckli, E. T. (2018). Understanding axon guidance: Are we nearly there yet? Development, 145 (10). 10.1242/dev.151415

Thompson, C. L., Ng, L., Menon, V., Martinez, S., Lee, C.-K., Glattfelder, K., Sunkin, S. M., Henry, A., Lau, C., Dang, C., Garcia-Lopez, R., Martinez-Ferre, A., Pombero, A., Rubenstein, J. L. R., Wakeman, W. B., Hohmann, J., Dee, N., Sodt, A. J., Young, R., … Jones, A. R. (2014). A high-resolution spatiotemporal atlas of gene expression of the developing mouse brain. Neuron, 83 (2), 309–323. 10.1016/j.neuron.2014.05.033

Turing, A. M. (1952). The chemical basis of morphogenesis [Publisher: The Royal Society]. Philosophical Transactions of the Royal Society of London. Series B, Biological Sciences, 237 (641), 37–72. Retrieved October 11, 2022, from https://www.jstor.org/stable/92463

Virtanen, P., Gommers, R., Oliphant, T. E., Haberland, M., Reddy, T., Cournapeau, D., Burovski, E., Peterson, P., Weckesser, W., Bright, J., Van Der Walt, S. J., Brett, M., Wilson, J., Millman, K. J., Mayorov, N., Nelson, A. R. J., Jones, E., Kern, R., Larson, E., … Vázquez-Baeza, Y. (2020). SciPy 1.0: Fundamental algorithms for scientific computing in python. Nature Methods, 17 (3), 261–272. 10.1038/s41592-019-0686-2

von der Malsburg, C. (1987). Ordered retinotectal projections and brain organization. In F. E. Yates, A. Garfinkel, D. O. Walter, & G. B. Yates (Eds.), Self-organizing systems: The emergence of order (pp. 265–277). Springer US. 10.1007/978-1-4613-0883-615

Waddington, C. H. (1956). The strategy of the genes. Routledge. 10.4324/9781315765471

Wolpert, L. (1969). Positional information and the spatial pattern of cellular differentiation. Journal of Theoretical Biology, 25 (1), 1–47. 10.1016/S0022-5193(69)80016-0

Zador, A. M. (2019). A critique of pure learning and what artificial neural networks can learn from animal brains. Nature Communications, 10 (1), 3770. 10.1038/s41467-019-11786-6

